# Neural representation of current and intended task sets during sequential judgements on human faces

**DOI:** 10.1101/753533

**Authors:** Paloma Díaz-Gutiérrez, Sam J. Gilbert, Juan E. Arco, Alberto Sobrado, María Ruz

## Abstract

Engaging in a demanding activity while holding in mind another task to be performed in the near future requires the maintenance of information about both the currently-active task set and the intended one. However, little is known about how the human brain implements such action plans. While some previous studies have examined the neural representation of current task sets and others have investigated delayed intentions, to date none has examined the representation of current and intended task sets within a single experimental paradigm. In this fMRI study, we examined the neural representation of current and intended task sets, employing sequential classification tasks on human faces. Multivariate decoding analyses showed that current task sets were represented in the orbitofrontal cortex (OFC) and fusiform gyrus (FG), while intended tasks could be decoded from lateral prefrontal cortex (lPFC). Importantly, a ventromedial region in PFC/OFC contained information about both current and delayed tasks, although cross-classification between the two types of information was not possible. These results help delineate the neural representations of current and intended task sets, and highlight the importance of ventromedial PFC/OFC for maintaining task-relevant information regardless of when it is needed.

## 1. Introduction

The selection and maintenance of relevant information is critical for our ability to pursue complex and hierarchically organized goals. In cases where we hold delayed intentions that need to be fulfilled later on (also known as prospective memory; Kliegel, McDaniel, & Einstein, 2008) or when we perform sequential tasks, it is necessary to represent the currently-active task and, in addition, the one to be performed later on. It is also important to switch flexibly from one task set to another (e.g. Monsell, 2003). Some studies have examined the neural representation of currently-active task sets in frontoparietal areas (e.g. Waskom, Kumaran, Gordon, Rissman, & Wagner, 2014; Woolgar, Thompson, Bor, & Duncan, 2011), while others have investigated the representations of delayed intentions suggesting a key role of medial prefrontal cortex (mPFC) in combination with more posterior areas (e.g. Gilbert, 2011; Haynes et al., 2007; Momennejad & Haynes, 2013). However, no previous study has examined the representation of current and intended task sets within a single experimental paradigm. This combination allows to investigate the extent to which currently-active and intended future task sets are represented in overlapping versus distinct brain networks, and also to contrast their activation patterns directly. Furthermore, previous studies have focused on representations of rather simple stimuli (i.e. geometric figures, objects, words, etc.; Crittenden, Mitchell, & Duncan, 2015; Waskom et al., 2014; Woolgar et al., 2011b), so it is not clear how well these findings generalize to more complex stimuli, such as human faces. In this study, we employed social categorization dual-sequential judgments on human faces to investigate the common and differential representation of current and delayed tasks.

The influence of maintaining an intended task-set on current task performance has previously been investigated with behavioural methods. These studies highlight how performance declines with an increment the number of tasks that need to be maintained, showing that the representation of two tasks simultaneously is more demanding compared to one task only. For instance, Smith (2003) found that participants performed an ongoing task more slowly when they held in mind a pending intention, compared with performing the ongoing task alone. This behavioural effect is accompanied by changes in pupil dilation (Moyes, Sari-Sarraf, & Gilbert, 2019), which also serves as an indicator of task demands (see van der Wel & van Steenbergen, 2018). Further, dual-task costs have also been manifested in task switching paradigms, where participants must switch between two active task-sets (Monsell, 2003; Rogers & Monsell, 1995). Even when the same task is repeated from the previous trial, responses are slower and less accurate during mixed blocks (where more than one task is relevant) compared with pure blocks consisting of just one task (Marí-Beffa, Cooper, & Houghton, 2012).

Results at the neural level also indicate that the maintenance of two tasks compared with one alters activity in specific brain regions. Several studies have shown that a set of “task-positive” regions increase their activation during demanding tasks (also known as the Multiple Demand network, MD; Duncan, 2010). This network is also sensitive to cognitive load, increasing its sustained activation as task complexity is raised (Dumontheil, Thompson, & Duncan, 2011; Palenciano, González-García, Arco, & Ruz, 2019; but see Tschentscher, Mitchell, & Duncan, 2017). Among these areas, the lateral prefrontal (lPFC) and parietal cortices play a prominent role during dual-task performance. Both increase their activation during task-switching trials while anterior PFC shows sustained activation during task-switching blocks (Braver, Reynolds, & Donaldson, 2003). Similarly, others (Szameitat, Schubert, Müller, & Von Yves Cramon, 2002) have shown involvement of lPFC during dual-task blocks, proportionally to task difficulty, during simultaneous and interfering task processing. Further, some studies have employed multivoxel pattern analysis (MVPA) to show how these frontoparietal (FP) regions code current task sets (Palenciano et al., *in press*; Qiao, Zhang, Chen, & Egner, 2017; Waskom et al., 2014; Woolgar, Hampshire, Thompson, & Duncan, 2011) and how the representation of task-relevant information in these areas increases with task demands (Woolgar et al., 2011b).

Traditionally, the role of FP regions has been opposed to “task-negative” areas, initially linked to decreased activity during effortful task performance (Fox et al., 2005), although recent studies suggest that it has a much broader role. This Default Mode Network (DMN) includes the ventro/dorsomedial PFC, orbitofrontal cortex (OFC), precuneus/posterior cingulate, inferior parietal lobe (IPL), lateral temporal cortex, and hippocampal formation (Buckner, Andrews-Hanna, & Schacter, 2008; Raichle, 2015). However, recent studies have qualified this view, showing that these regions also represent task-relevant information in different contexts (e.g. Crittenden et al., 2015; González-García et al., 2017; Palenciano et al., *in press*; Smith, Mitchell, & Duncan, 2018). Moreover, functional connectivity approaches have shown that the strength of connectivity among task-negative regions during a working memory task is associated with better performance (Hampson, Driesen, Skudlarski, Gore, & Constable, 2006). Similarly, Elton and Gao (2015) observed that the dynamics of connectivity among DMN regions during task performance were also related to behavioural efficiency. Altogether, the literature suggests a clear involvement of FP areas in the representation of current task-related information and highly demanding tasks. Conversely, the role of the DMN is less clear. Although it shows decreased activation during demanding tasks, its dynamics are also related to behaviour, and contain task information in different contexts. This suggests that these regions play a role in the representation of task-relevant knowledge.

Further, one of the main nodes of the DMN, the medial prefrontal cortex (mPFC) has an important role in the representation of intended behaviour during both task-free situations (Haynes et al., 2007) and delays concurrent with an ongoing task (Gilbert, 2011; Momennejad & Haynes, 2012; Momennejad & Haynes, 2013). This area also plays a role when holding decisions before they reach consciousness (Soon, Brass, Heinze, & Haynes, 2008). The evidence from studies of delayed intentions has led to suggested dissociations between the role of lateral and medial PFC (associated with the task-positive and task-negative networks, respectively). Momennejad & Haynes (2013) directly compared the representation of future intentions during delays with and without an ongoing task, and found that while the lPFC had a general role of encoding intentions regardless of whether there was or not an ongoing task during the delay, the mPFC was involved when the delay period was occupied by an ongoing task. Alternatively, Gilbert (2011) could not find encoding of delayed intentions in the lPFC but they did in the mPFC, suggesting that the former may play a content-free role in remembering delayed intentions while the latter would represent their specific content. However, these studies vary in the abstraction of the task rules employed. While Gilbert (2011) aimed to decode specific visual cues and responses, others focused on the anticipation of abstract task sets, such as arithmetic operations (addition vs. substraction; Haynes et al., 2007), or parity vs. magnitude judgements (Momennejad & Haynes, 2012; Momennejad & Haynes, 2013). This difference in abstraction could impact the brain region (lPFC vs. mPFC) maintaining information about future intentions (Momennejad & Haynes, 2013). Further, these studies also vary in whether the retrieval of the intended task was cued (Gilbert, 2011) or self-initiated (Momennejad & Haynes, 2012; Momennejad & Haynes, 2013). Therefore, although these studies have studied the representation of intentions in a variety of experimental settings, they have not directly addressed how the representation of a future task set may differ from the representation of an ongoing task that is currently being performed.

In addition, the studies so far have employed mainly non-social stimuli. In this context it is worth noting that the DMN has also been related to processes relevant in the social domain (Buckner & Carroll, 2007; Mars et al., 2012; Spreng, Mar, & Kim, 2008). For instance, engagement of the DMN during rest is related to better memory for social information (Meyer, Davachi, Ochsner, & Lieberman, 2018). Facial stimuli are an important source of social knowledge, which is represented in a set of regions including the fusiform gyri (FG; Haxby, Hoffman, & Gobbini, 2000; Kanwisher & Yovel, 2006). This FG also shows different neural patterns distinguishing social categories (Kaul, Ratner, & Van Bavel, 2014; Stolier & Freeman, 2017). Similarly, the representational structure of social categories is altered by personal stereotypes both in the FG and in higher-level areas such as the OFC (Stolier & Freeman, 2016), which is also linked to the representation of social categories such as gender, race, or social status (Gilbert, Swencionis, & Amodio, 2012; Kaul, Rees, & Ishai, 2011; Koski, Collins, Olson, & Hospital, 2017) and the integration of contextual knowledge during face categorization (Freeman et al., 2015). Likewise, during predictive face perception, the FG coactivates with and receives top-down influences from dorsal and ventral mPFC (e.g. Summerfield et al., 2006), which in turn have also been implicated on judgements about faces (Mitchell, Macrae, & Banaji, 2006; Singer, Kiebel, Winston, Dolan, & Frith, 2004). Therefore, given the special properties and influence of social information gathered from faces, understanding how task-relevant current and delayed information may be represented when it pertains to social information is important to extend and complement previous findings.

In the current fMRI study, we employed a dual-sequential categorization task, where participants had to discriminate between features of three dimensions of facial stimuli and had to maintain for a period of time both the initial ongoing task and an intended one. In particular, we studied how demands (one vs. two sequential tasks) influence performance, and hypothesized that high demand would be associated with worse performance alongside with activation in frontoparietal regions, especially the lPFC. To examine the brain regions containing fine-grained information about both current and intended tasks we employed MVPA. Unlike traditional univariate methods, where the mean activation in a set of voxels is compared between conditions, MVPA focuses on the spatial distribution of activations. Here, a classifier is trained to distinguish response patterns associated with different experimental conditions (i.e. stimuli categories, cognitive states, etc.) in a certain brain region. If the trained classifier is able to predict the patterns of independent data, there is indication that the brain area under study represents specific information about those conditions. Thus, MVPA allows to examine finer-grained differences in how information is represented in the brain (for reviews see Haxby, Connolly, & Guntupalli, 2014; Haynes, 2015). In this work, we aimed to study how an intended task set might be represented differently from a currently-active ongoing task. For that reason, we focused on the initial pre-switch period, when the current task is being performed before switching to the intended task. Specifically, we performed separate analyses to decode: 1) the task currently being performed, regardless of the intended future task; 2) the task intended for the future, regardless of the current task (henceforth: “initial task” and “intended task”, respectively). Given the extensive literature associating FP areas to the representation of task-relevant information (Qiao et al., 2017; Waskom et al., 2014; Woolgar et al., 2011a, 2011b), we expected to decode the initial relevant task in MD regions and the intended one in “task-negative” regions, especially the mPFC, in line with previous studies showing its role in prospective memory (Gilbert, 2011; Haynes et al., 2007; Momennejad & Haynes, 2012; Momennejad & Haynes, 2013).

## 2. Methods

### 2.1. Participants

Thirty-two volunteers were recruited through adverts addressed to undergraduates and postgraduate students of the University of Granada (range: 18-28, M = 22.5, SD = 2.84, 12 men). All of them were Caucasian, right-handed with normal or corrected-to-normal vision and received economic remuneration (20-25 Euros, according to performance) in exchange for their participation. Participants signed a consent form approved by the Ethics Committee for Human Research of the University of Granada.

### 2.2. Apparatus and stimuli

We employed 24 face photographs (12 identities, 6 females, 6 black; 3 different identities per sex and race) displaying happy or angry emotional expressions, extracted from the NimStim dataset (Tottenham et al., 2009). E-Prime 2.0 software (Schneider, Eschman, & Zuccolotto, 2002) was used to control and present the stimuli on a screen reflected on a coil-mounted mirror inside the scanner.

### 2.3. Design and procedure

Participants had to perform a series of categorization tasks where they judged either the emotion (happy vs. angry), the gender (female vs. male) or the race (black vs. white) of series of facial displays. These tasks were arranged in miniblocks, which could each contain one (Pure Miniblock; PM) or two sequential categorization tasks (Mixed Miniblock; MM). At the beginning of each miniblock, participants received instructions indicating the number of tasks to perform (1 vs. 2) and their order and nature (Emotion, Gender and/or Race), as well as the key-response mappings. Thus, for PMs, the initial instruction indicated the one task that had to be performed during the whole miniblock. Conversely, for MMs, the instruction indicated two tasks, where the participant had to change from the first to the second task at a certain point of the miniblock. After the instruction, a coloured (blue or red) fixation point appeared on the screen, followed by a facial display (see Figure 1). Participants were told that, during MMs, they had to switch tasks when the fixation changed its colour (from blue to red or vice versa). Once it switched, they had to continue doing the second task until the end of the miniblock. To equate the perceptual conditions across blocks, the fixation colour also changed during PMs, although participants were told to ignore this change.

**Figure 1.**
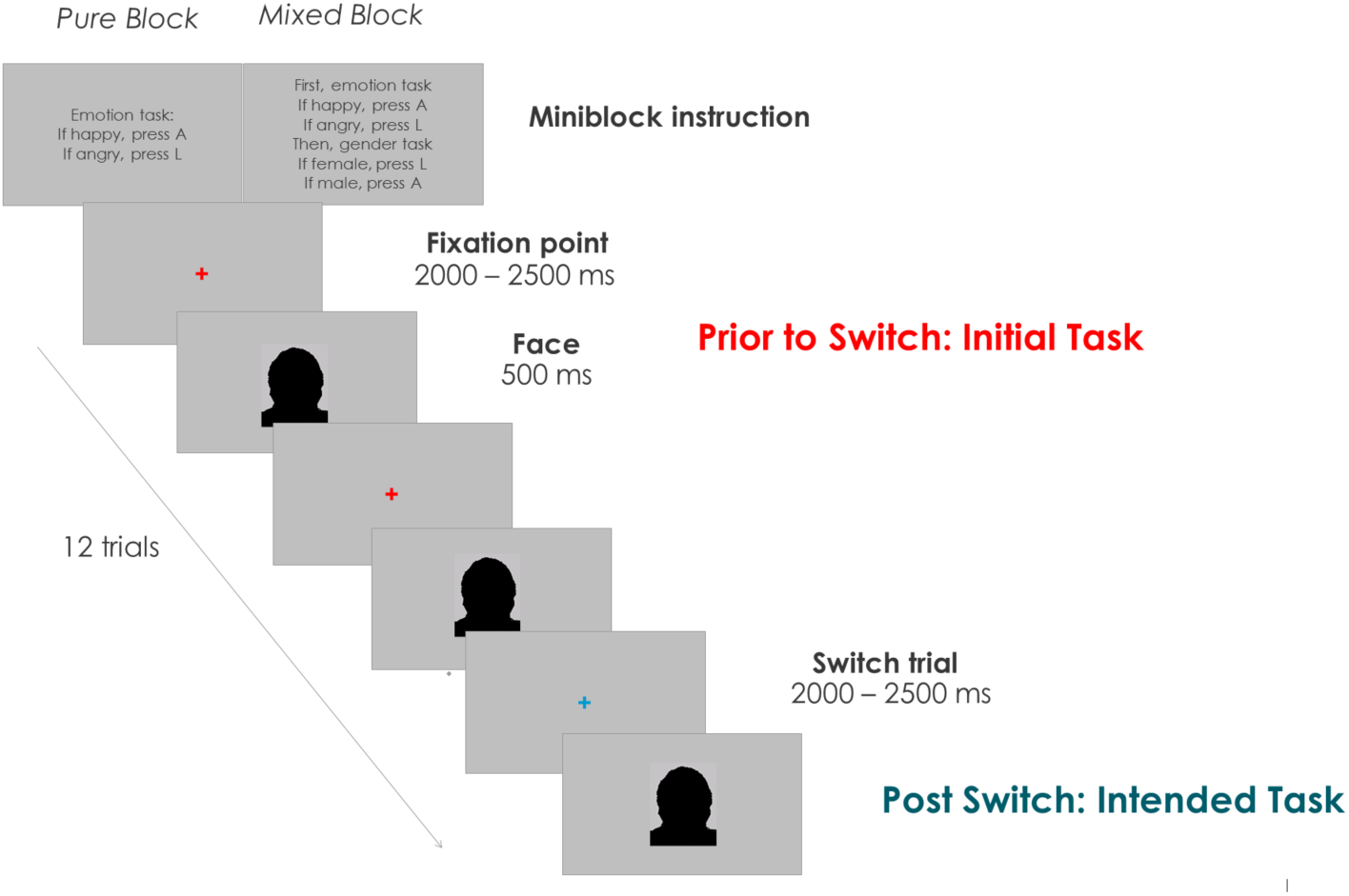
Display of the paradigm (face stimuli are obscured in the preprint version). Example of a miniblock and sequence of trials. Inter-trial-interval (ITI) duration = 2-2.5 s.

Hence, in each MM there was an initial task (first task to perform), an intended task (second task to perform) and an ignored task (non-relevant for that miniblock). Importantly, our main fMRI analyses focused on the period before the switch, while participants needed to represent both the initial task and the intended one. Task switches were evenly spaced across the miniblock, from trial 1 to 12. This allowed us to decorrelate brain activity associated with the pre-switch period, post-switch period, and the switch itself. In total, there were 9 different types of miniblocks: 3 pure (emotion[EE], gender[GG], race[RR]) and 6 mixed (emotion-gender[EG], emotion-race[ER], gender-emotion[GE], gender-race[GR], race-emotion[RE], race-gender[RG], see Figure 2). Across the experiment, pure miniblocks appeared 8 times each, while every type of mixed miniblock was repeated 12 times. The presentation order of the miniblocks and the assignment of response options (left or right index) were counterbalanced within each run. Additionally, to avoid response confounds in the analyses, response mappings changed between runs. Thus, for each participant odd and even runs had the opposite response mappings.

**Figure 2.**
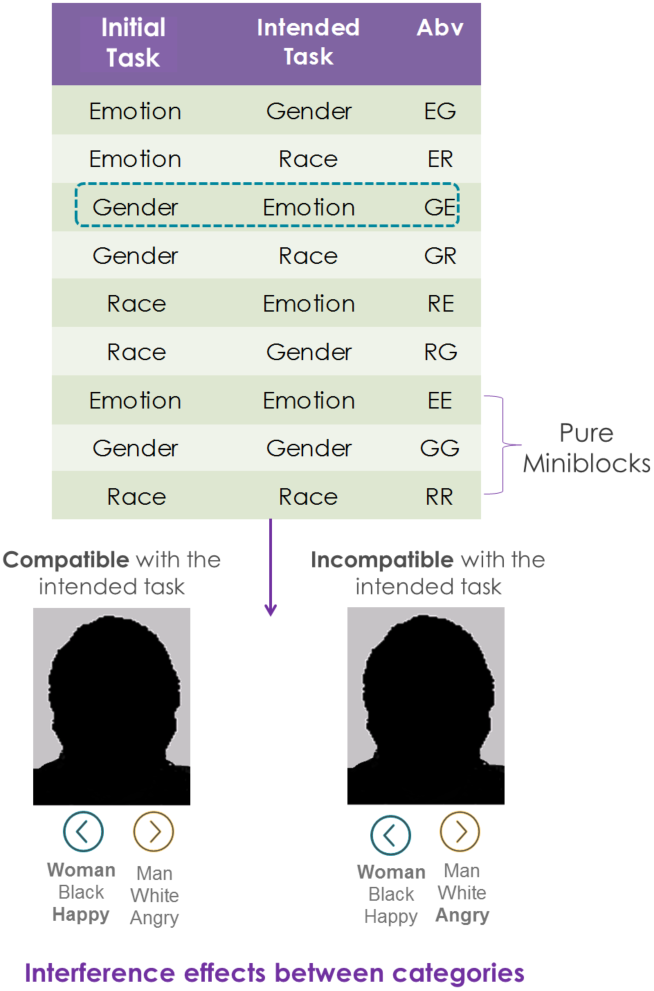
Top: All possible combinations of miniblocks, depending on the initial and intended tasks, and their abbreviation (Abv). Bottom: Example of interference between initial and intended categories in a Gender-Emotion (GE) miniblock.

Participants performed a practice block to learn the different tasks and the response mappings. They were required to obtain a minimum of 80% accuracy at this practice block prior to entering the scanner. The sequence of each miniblock was as follows: First, the instruction slide presented the task/s to perform (Pure: 1, Mixed: 2), and the response mappings (right/left index), during 5 s. Then, a sequence of 12 trials appeared. In each of them, a fixation point (blue or red, counterbalanced) lasting 2-2.5 s (inter-trial-interval; ITI; in units of 0.25 s, randomly assigned to each fixation) was followed by a facial display of 0.5 s. The fixation for the switch trial lasted on average 2.24 s (SD = 0.022; all participants within a range of ± 2.5 standard deviations). The experiment consisted of 1152 trials, arranged in 96 miniblocks (72 mixed and 24 pure), distributed in 12 scanning runs. Hence, each run consisted of 8 miniblocks (6 mixed and 2 pure). Each type of miniblock was repeated 12 times, once per run, and each time the switch occurred on a different trial. Presentation order and switch point were counterbalanced through the experiment, to ensure that each switch occurred on every possible trial for each type of miniblock and that each identity was associated the same number of times with the switch. In total, the fMRI task lasted for 60.8 min.

In addition, we also studied the interference between tasks. Since in MMs the participant had to perform two tasks sequentially, the established stimulus-response association could be compatible or incompatible between the current and intended task, depending on the specific target face. For instance, the gender task could have a stimulus-response (S-R) association of female-left/male-right and the S-R in the emotion task might be happy-left/angry-right. Therefore, in a GE miniblock, during the gender task, participants could encounter happy female faces (both the initial gender and intended emotion tasks would require the same response: left) or angry female faces (the initial gender task would lead to response with the left index and the emotion with the right one). Thus, the former would be an example of compatibility between initial and intended tasks, whereas the latter would entail incompatibility (see Figure 2).

### 2.4. Image acquisition and preprocessing

Volunteers were scanned with a 3T Siemens Magnetom Trio, located at the Mind, Brain and Behavior Research Center (CIMCYC) in Granada, Spain. Functional images were obtained with a T2*-weighted echo planar imaging (EPI) sequence, with a TR of 2.210 s. Forty descending slices with a thickness of 2.3 mm (20% gap) were extracted (TE = 23 ms, flip angle = 70 °, voxel size of 3×3×2.3 mm). The sequence was divided into 12 runs, consisting of 152 volumes each. Afterwards, an anatomical image for each participant was acquired using a T1-weighted sequence (TR = 2500 ms; TE = 3.69 ms; flip angle = 7°, voxel size of 1 mm^3^). MRI images were preprocessed and analysed with SPM12 software (http://www.fil.ion.ucl.ac.uk/spm/software/spm12). The first 3 images of each run were discarded to allow the stabilization of the signal. The volumes were realigned and unwarped and slice-time corrected. Then, the realigned functional images were coregistered with the anatomical image and were normalized to 3 mm^3^ voxels using the parameters from the segmentation of the anatomical image. Last, images were smoothed using an 8 mm Gaussian kernel, and a 128 high-pass filter was employed to remove low-frequency artefacts. Multivariate analyses used non-normalized and non-smoothed data (Bode & Haynes, 2009; Gilbert & Fung, 2018; Woolgar et al., 2011a, 2011b).

### 2.5. fMRI analyses

#### 2.5.1. Univariate

First, we employed a univariate approach to examine the effect of context demands (one vs. two sequential tasks) and task switching. Our model contained, for each run, one regressor for the instruction of each miniblock, four regressors corresponding to the two types of miniblock (pure/mixed) with separate regressors for the pre-switch and post-switch periods, one for the change in fixation colour during mixed miniblocks (indicating a switch event), one for the change in fixation colour during pure miniblocks (serving as a baseline for the switch events), and another one for the errors. Both instruction and miniblock regressors were modelled as a boxcar function with the duration of the entire pre/post switch period or instruction duration (5 s). Errors were modelled including the duration of the face of that trial and the following fixation (2.5-3 s). Switch trials were modelled as events, with stick functions with zero duration locked at the switch in colour of the fixation point. This provided a model with a total of 8 regressors per run. At the group level, t-tests were carried out for comparisons related to the effect of task demands (one vs. two tasks) at the period prior to the switch, and also to compare switching cost effects (switch trial in the mixed block > switch trial in pure blocks). We report clusters surviving a family-wise error (FWE) cluster-level correction at p< .05 (from an initial uncorrected threshold p < .001). Additionally, we also performed nonparametric inference (see Supplementary Materials).

#### 2.5.2. Multivariate analysis

We performed multivoxel pattern analyses (MVPA) to examine the brain areas maintaining the representation of A) current-active initial tasks, and B) intended tasks. These analyses examined brain activity during the pre-switch period only (although additional, exploratory analyses were also performed on the post-switch period, see the Supplementary Materials). Following a Least-Squares Separate Model approach (LSS; Turner, 2010) we modelled each miniblock (EG, ER, GE, GR, RE, RG) during the period prior to the switch separately. This method helps to reduce collinearity between regressors (Abdulrahman & Henson, 2016), by fitting the standard hemodynamic response to two regressors: one for the current event (a type of miniblock prior to the switch) and the second one for all the remaining events. As in the univariate approach, each miniblock regressor was modelled as a boxcar function with the duration of the entire pre-switch period duration.

The binary classification analyses were performed as follows. First, we classified a) between any two initial tasks while holding the intended task constant, then b) between any two intended tasks while holding the initial task constant. For instance, in case a) we contrasted initial gender task vs. initial race task when the intended task was emotion (GE vs. RE), and also intended gender vs. intended emotion task when the intended task was race (ER vs. GR), and intended emotion task vs. intended race task when the intended task was gender (EG vs. RG). We then averaged decoding accuracies across these analyses, which indicate whether a particular brain region shows different patterns of activity depending on what the initial, currently-active task set is, holding the intended task constant. Conversely, in case b), we compared intended gender vs. intended race when the initial, currently-active task was emotion (EG vs. ER), intended gender vs. intended emotion when the initial, currently-active task was race categorization (RE vs. RG) and intended emotion vs. intended race when the initial, currently-active task was gender (GE vs. GR). As above, we averaged across these analyses, which indicate whether a particular brain region shows different patterns of activity depending on what the intended task set is, holding the currently-performed task constant.

To carry out these analyses, we performed a whole brain searchlight (Kriegeskorte, Goebel, & Bandettini, 2006) on the realigned images employing the Decoding Toolbox (TDT; Hebart, Görgen, & Haynes, 2015) and custom-written MATLAB code. We created 4-voxel radius spheres and for each sphere, a linear support vector machine classifier (C =1; Pereira, Mitchell, & Botvinick, 2009) was trained and tested using a leave-one-out cross-validation. Due to the nature of the paradigm and the counterbalancing, once in each block the switch took place at the first trial (here particcipants only performed the intended task). Thus, there was an example of each type of miniblock before the switch in only 11 runs, differently for each participant and miniblock (i.e. a participant could lack miniblock EG in run 4 and miniblock RG in run 11). To avoid potential biases in the classifier for having only one of the classes in a run, for each participant and comparison, we performed the classification only in the 10 runs where there was an example of both miniblocks. Resulting from this procedure, we employed the data from 10 scanning runs (training was performed with data from 9 runs and tested on the remaining run, in an iterative fashion). In the exceptional case (twice for each contrast in the total sample) where the two miniblocks in the classification were absent in the same run, classification was performed on the remaining 11 runs (training with data from 10 runs and testing on the remaining run). In addition, we observed biases in the decoding estimates when the switch trial from one of the conditions in the test set matched the opposite class in the training set, which happened for every comparison in aproximately half of the cross-validation steps. To avoid the biases resulting from this, we additionally removed those runs where the switch position matched the test from the training set for that specific cross-validation step.

Next, we averaged the accuracy maps for a) and b) to obtain a mean classification map collapsing across initial and intended tasks. This allowed us to detect regions that contained information about either initial or intended tasks (or both). It also allowed us to define ROIs that could be used to compare decoding accuracies for initial versus intended task-sets, in a manner that was unbiased between the two types of information. We additionally conducted whole-brain analyses investigating decoding of the initial task only, decoding of the intended task only, and the comparisons between the two.

Afterwards, group analyses were performed by doing one-sample t-tests after normalising (same as for the univariate analyses) and smoothing the individual accuracy maps (4 mm Gaussian kernel, consistent with earlier MVPA studies such as Gilbert, 2011; Gilbert & Fung, 2018). Results were considered significant if they passed an FWE cluster-level correction at p< .05 (based on an uncorrected forming threshold of p< .001). This statistical approach is consistent with recent MVPA studies (Gilbert & Fung, 2018; Loose, Wisniewski, Rusconi, Goschke, & Haynes, 2017). We additionally carried out nonparametric inference (see Supplementary Materials).

## 3. Results

### 3.1. Behaviour

First, to study how the number of tasks influenced performance, we performed a paired t-test on both accuracy and reaction times (RTs), between the two types of Miniblock (Mixed/Pure), collapsing over pre- and post-switch periods (see Figure 3). Here, responses were more accurate for pure (M = 95.7%, SD = 4.3), than for mixed miniblocks (M = 92.3%, SD = 3.6), t_31_ = 5.39, p<.001, whereas they were faster for pure (M = 671.12 ms, SD = 126.3) than for mixed (M = 709.18 ms, SD = 142.39) miniblocks, t_31_ = 4.83, p<.001.

**Figure 3.**
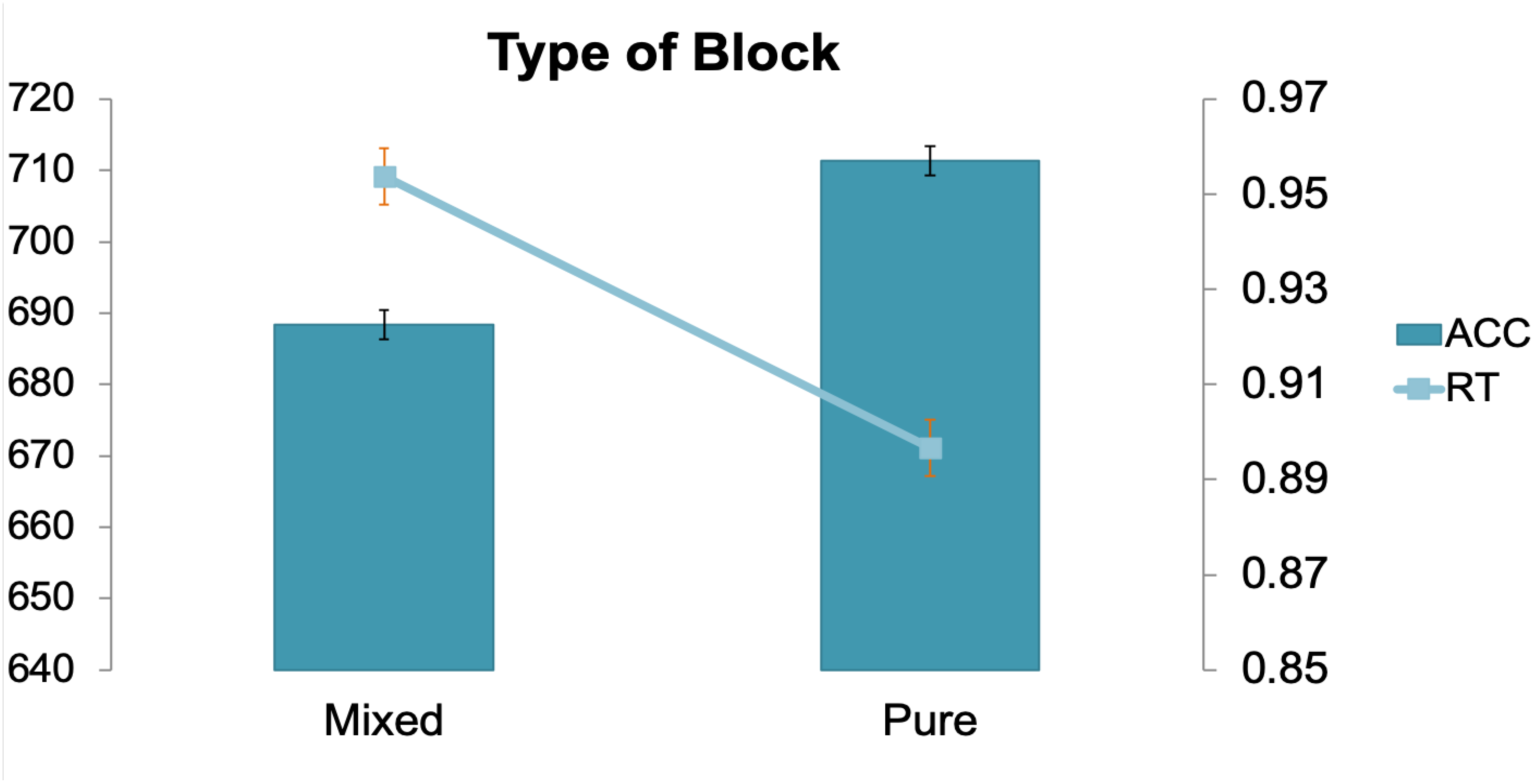
Influence of the type of block on performance. Error bars represent within-subjects 95% confidence intervals (Cousineau, 2005).

In addition, we examined if the intended task influenced initial task performance, and vice versa. For this, we selected only the mixed miniblocks and entered them into a repeated measures (rm) ANOVA, with Task (Emotion/Gender/Race), Period of the miniblock (Prior to/Post Switch), and Interference (Compatible/Incompatible) between initial and intended tasks. Note here that even if we did not have any specific hypothesis about the influence of the variable Task on performance, we included it as a factor in this second ANOVA to examine whether the Task modulated the effect of the other two variables of interest: Period of the block and Interference. Moreover, we refer to initial and intended as the tasks performed before and after the switch, respectively, to preserve consistency in the terminology throughout the entire manuscript. In addition, we refer as compatible trials when the correct response for the initial task-relevant dimension was associated with the same response for the intended dimension (see an example in Figure 1, right), and incompatible trials when the response associated with the initial task-relevant dimension interfered with the responses associated with the intended one. Similarly, after the switch, when the intended task was being performed, compatible trials referred to those where the correct responses for this task were associated with the same response for the previous initial task, and incompatible trials when the response associated with both dimensions differed. This way we could use interference effects as an indicator of the maintenance of the intended response dimensions during performance.

Accuracy did not show any main effect of Task (F<1, see Figure 4). However, we observed a main effect of Period of the miniblock, F_1,31_ = 58.215, p < .001, ηp^2^ = .653, where participants responded more accurately before (M = 94.86%, SD = 3.6) than after the task switch (M = 90.05%, SD = 5.4). There was also a main effect of Interference, F_1,31_ = 101.83, p < .001, ηp^2^ = .767, where accuracy was higher for compatible (M = 94.91%, SD = 4.76) than for incompatible trials (M = 90%, SD = 6.79). The interaction Task x Period was significant, F2,62 = 3.831, p = .029, ηp^2^ = .110, showing that performance was better before than after the switch for all three tasks (all Fs > 15, ps < .001), but this difference was larger for the gender task (ηp^2^ = .679). Similarly, the interaction of Period x Interference was significant, F_1,31_ = 52.244, p < .001, ηp^2^ = .106, where accuracy was worse for incompatible compared to compatible trials (both Fs > 19, ps < .001), but this pattern was more pronounced after (F_1,31_ = 107.08, p<.001) than before the switch (F_1,31_ = 19.21, p<.001). No other interactions reached significance (p>.061).

**Figure 4.**
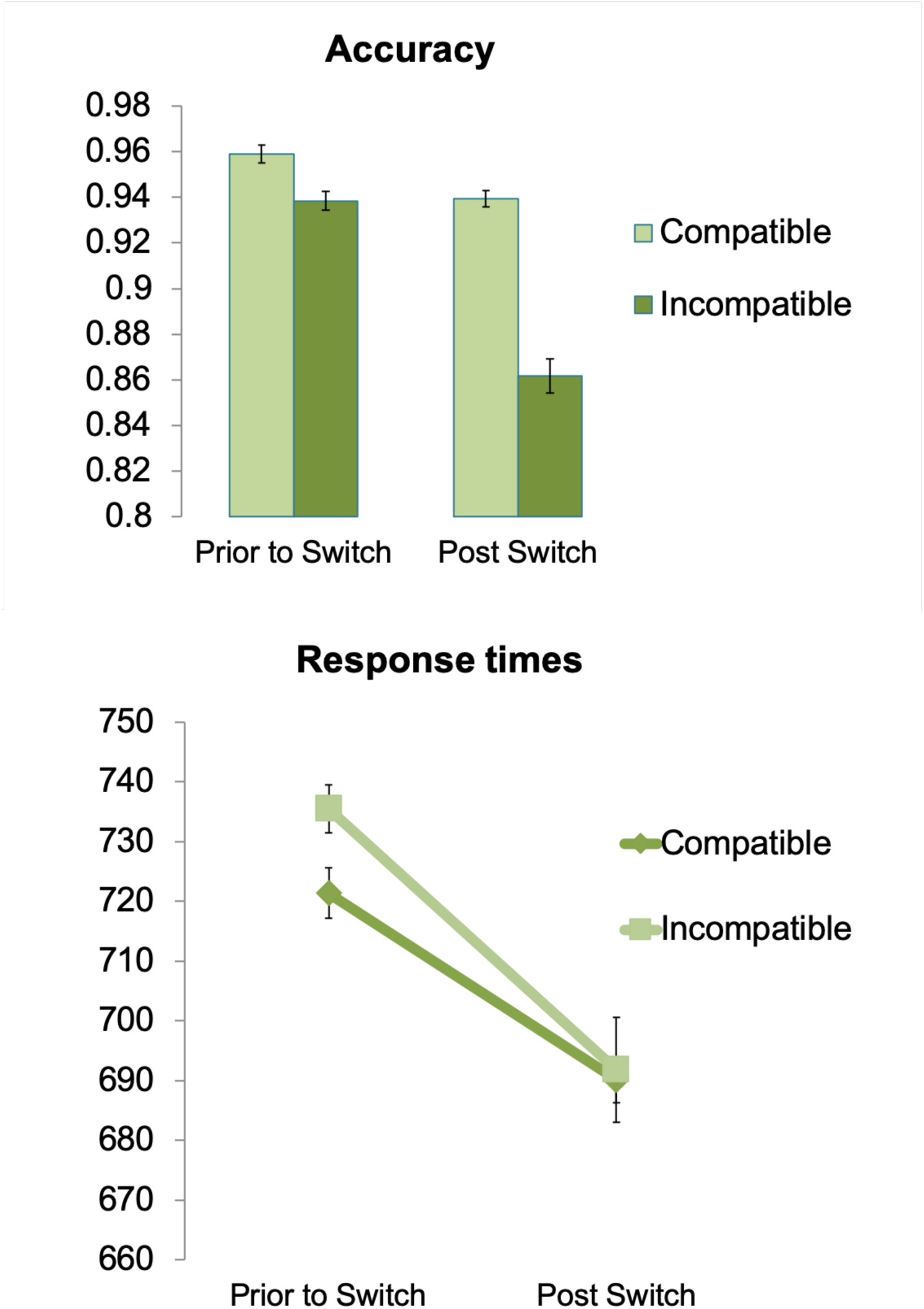
Interference effects between initial and intended tasks before and after the switch (MMs). Top: Accuracy rates. Bottom: Reaction times (ms). Error bars represent within-subjects 95% confidence intervals (Cousineau, 2005).

**Figure 5.**
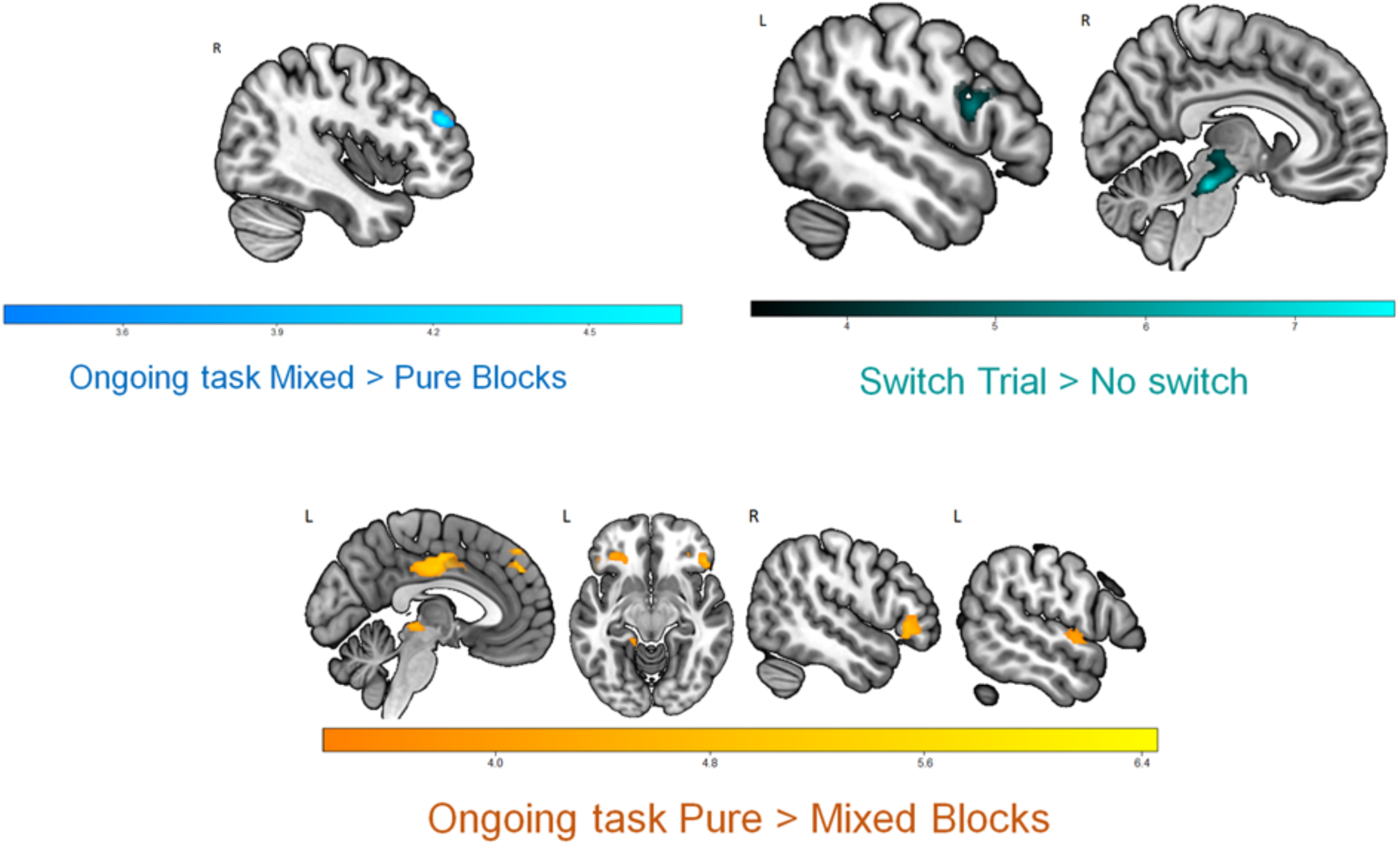
Univariate results. Effect of task demands (one vs. two) and task switching. Scales reflect peaks of significant t-values (p<.05, FWE-corrected for multiple comparisons).

**Figure 6.**
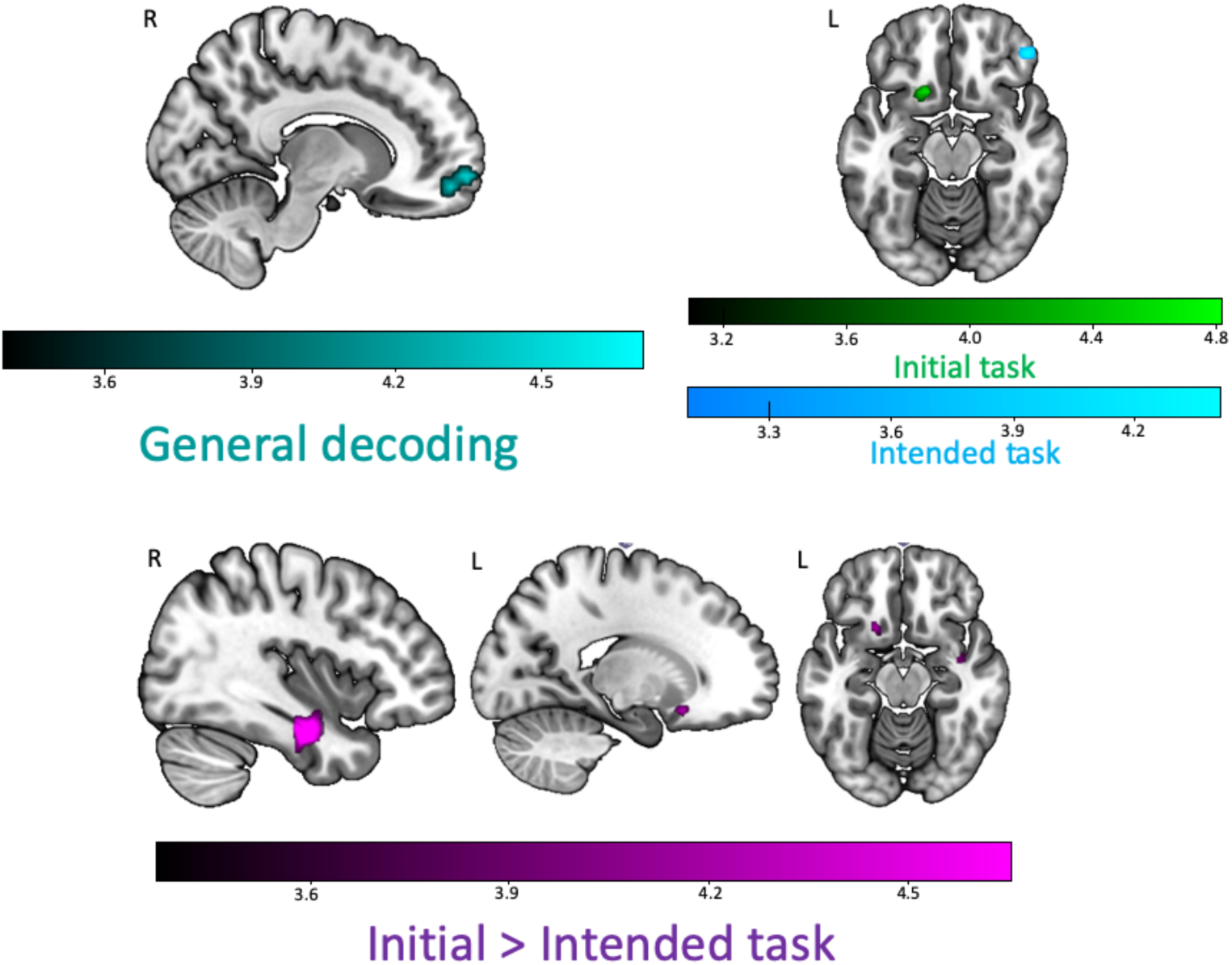
Multivariate results during the period prior to the switch. Top left: General decoding (cyan) of the region sensitive to any kind of task (initial or intended). Top right: Decoding of the initial (green) and intended (blue) task separately. Bottom: Decoding of initial > intended task (violet). Scales reflect peaks of significant t-values (p<.05, FWE-corrected for multiple comparisons).

RTs showed (see Figure 4) a main effect for Task (F2,62 = 24.08, p < .001, ηp^2^ = .437), were race was performed faster (M = 691.16, SD = 150.07), followed by gender (M = 709.29, SD = 147.11), and emotion (M = 728.59, SD = 144.14). In addition, we also observed a main effect of Period (F_1,31_ = 32.83, p < .001, ηp^2^ = .514), as participants were faster after (M = 690.94, SD = 143.38) than prior to the switch (M = 728.42, SD = 155.72). Further, we found a main effect of Interference, F_1,31_ = 4.829, p =.036), where participants were faster for compatible (M = 705.51, SD = 147.79) than incompatible trials (M = 713.65, SD = 146.42). An interaction Period x Interference (F1,32 = 24.08, p = .033, ηp^2^ = .437) showed that this interference effect was significant before the switch (F_1,31_ = 8.15, p = .008), but not after (F_1,31_ = .178, p>.67). None of the other interactions were significant (p>.2).

During mixed blocks we observed higher accuracies and reaction times during the period before the switch and the opposite pattern (low accuracies and faster responses) after it, which could indicate a trade-off in the data. To address this possibility, we additionally performed Pearson correlations between mean accuracy and reaction times during both periods of the miniblock. Results show no association between the two measures, neither before (r = .07; p > .35) or after (r = .17, p > .17) the switch.

### 3.2. fMRI

#### 3.2.1. Univariate

##### 3.2.1.1. Pure vs. Mixed blocks

Before the switch, the right middle frontal gyrus (k = 56; MNI coordinates of peak voxel: 42, 44, 23) showed higher activation when participants had to maintain two tasks vs. one (Mixed > Pure blocks). Conversely, in this scenario, we observed decreased activation (Pure > Mixed blocks) in a set of regions. These included the bilateral middle cingulate cortex (k = 251; −3, −10, 33), bilateral medial prefrontal cortex (mPFC; k = 80; −6, 44, 41), left orbitofrontal cortex (OFC; k = 117; −33, 32, −16), right inferior frontal gyrus (IFG)/OFC (k = 168; 54, 32, −1), left lingual and parahippocampal gyri (k = 148; −9, 52, 5) and left superior temporal gyrus (k = 62; −57, −1, −4).

##### 3.2.1.2. Switch vs. non-switch trials

Transient activity during task switching was observed in the bilateral brainstem and thalamus (k = 291; −3, −28, −22) as well as in a cluster including the left inferior/middle frontal gyrus (IFG/MFG) and precentral gyrus (k = 160; −51, 11, 17).

##### 3.2.1.3. Interference effects

We assessed whether the interference effects observed on behaviour were matched at a neural level, by comparing incompatible vs. compatible trials before and after the switch. For this model, we included in each run one regressor for the instruction of each miniblock, one regressor corresponding to all Pure Miniblocks, four regressors corresponding to Compatible/Incompatible trials in Mixed Miniblocks, with separate regressors for the pre-switch and post-switch periods (onset at face presentation) and a last regressor for errors. Instructions and errors were modelled as described in section

#### 2.5.1 Univariate analysis

Pure Miniblocks were modelled as a boxcar function with the duration of the entire pre/post switch period, while the four remaining regressors (Compatible Pre, Compatible Post, Incompatible Pre, Incompatible Post) were modelled as events with zero duration. This provided a model with a total of 7 regressors per run. At the group level, t-tests were carried out for comparisons related to the effect of Interference before and after the switch. No effect survived multiple comparisons. However, at a more lenient threshold (p<.001, uncorrected), data showed higher activation in the left IFG (k = 22; −48, 35, 20) for incompatible > compatible trials. The opposite comparison (compatible > incompatible before the switch) or incompatible vs. compatible contrasts after the switch did not yield any significant results.

#### 3.2.2. Decoding results

First, we averaged all individual classification maps to examine the regions sensitive to any kind of task (initial or intended) during the period prior to the switch. We found that the rostromedial PFC/orbitofrontal cortex (OFC) presented significant accuracies above chance (k = 68; 15, 56, −10). To further examine whether one of the tasks dominated the classification, we extracted the decoding values from the initial and intended classification from the general decoding ROI above. A paired t-test between the initial and intended task decoding values showed no differences between decoding accuracy for the two types of information, t_31_ = .846, p = .404. To further test this idea, we ran an ROI analysis in this region to see whether decoding accuracy was significant for both the initial and intended task. To avoid non-independency, we employed a Leave-One-Subject-Out (LOSO) cross-validation approach (Esterman & Yantis, 2010) to select the ROI per participant. That is, data from each participant was extracted from an ROI that was defined based on the data from all the other participants, to avoid ‘double dipping’ (Kriegeskorte, Simmons, Bellgowan, & Baker, 2009). After this, both the initial and intended decoding values showed significant accuracy above chance (1-tailed, *t*31 = 3.018, *p*=.0025 and *t*31 = 2.299, *p* = .0145, respectively). This suggests that the mPFC/OFC region, prior to the switch, carries information about the relevant tasks to perform, regardless of whether they are initial or intended.

To further characterise the information represented in the mPFC/OFC region, we additionally examined whether we could cross-classify between the initial and intended tasks, which speaks to the potential overlap between these representations. Therefore, we performed ROI cross-classification analysis (Kaplan, Man, & Greening, 2015) in the mPFC/OFC region from the general decoding. We followed the same classification procedure as described in the methods section, but training the classifier on the initial task and testing on the intended task, and vice versa. Note that in this scenario the cross-classification was carried out in a totally independent and orthogonal analysis to that employed to define the OFC region. However, results showed no significant cross-classification in this region, t_31_ = 1, p > .3, which suggests that the initial vs. intended nature of the tasks may change their representational format.

Moreover, we examined the initial and intended individual classification maps separately to examine the regions sensitive to each type of task (initial or intended). With this, the classification of the initial task alone showed a different cluster in the left OFC (k = 52, −15, 17, −13). Conversely, we observed the right IFG (k = 53, 45, 41, −16) for the classification of the intended task alone. Last, when comparing decoding accuracies between the initial and intended tasks (subtracting initial – intended accuracy maps), we observed significantly higher accuracies for the representation of the initial (vs. intended) in the right FG and the hippocampus (k = 148; 39, −13, −25) and also in left OFC (k = 44; −18, 17, −16). However, the opposite contrast (intended vs. initial) did not show any cluster with significant differences.

Importantly, these MVPA results were unlikely to be due to differences in response times (see Supplementary Materials *“RT analyses between miniblock pairs and correlation with decoding accuracies”*).

Last, although the main focus of this work is the period before the switch, we performed exploratory classification analysis in the period after the switch to obtain further understanding about how information is represented once the initial task is no longer active. Following the same procedure as in section *2.5.2* *Multivariate analysis*, we carried out the decoding analysis with regressors of interest corresponding to the period after the switch. As in the previous analyses, we first averaged all individual classification maps to examine the regions sensitive to any kind of task during the period after the switch. Here, we observed a cluster in left middle frontal/precental gyrus (*k* = 61; −36, 11, 35) with significant accuracies above chance. Next, we averaged the classification maps separately for the currently-active post-switch task and the previously-performed initial task, to examine the regions sensitive to each type of task. The initial task could be decoded from bilateral IFG (left; *k* = 48; −54, 29, 26/right; *k* = 45; 54, 32, 5). Conversely, decoding of the currently-active post-switch task could only be observed at a more liberal threshold (p< .01, k = 100) in a cluster including the FG/cerebellum (*k* = 109; 15, −52, −19) and in left middle/inferior frontal gyrus (*k* = 122; −45, 8, 35).

## 4. Discussion

The present work aimed to examine the representation of A) currently-active, initial task sets, and B) intended task sets applied to social stimuli during a dual-sequential social categorization of faces. We found some regions from which we could decode only the currently-active or the intended task set, along with a region of vmPFC/OFC containing both types of information although cross-classification between the two was not possible.

The paradigm employed revealed behavioural costs due to the maintenance of intended task sets. Here, mixed blocks presented slower responses and lower accuracies, compared to pure ones, in line with previous studies (Los, 1996; Marí-Beffa et al., 2012). Response times were slower before than after the switch, when the intended task needed to be held while performing the initial one, in line with Smith (2003). These results reflect the higher demands associated with the maintenance of two relevant task sets. In addition, people need more time to respond when they hold the intention while performing the initial task, but not after, when they only need to focus on a single task. Further, we aimed to examine the activation of both initial and intended task sets before the switch by looking at possible interference between them. Behavioural results showed incompatibility effects in performance when participants needed to hold the intended task set while performing the ongoing task. These results point to the maintenance of intended task settings, showing that information about the pending set is maintained online during performance of the ongoing task.

Turning to brain activation data, the behavioural costs we observed during the maintenance of two task sets during mixed blocks were accompanied by increased activation in the right MFG, while regions such as the left IFG and the thalamus increased their activation during switching trials. Incompatible trials during this period were also related to activation in left lPFC, but only at an uncorrected statistical threshold. These results fit with previous work that associated lPFC with sustained control during dual tasks (Braver et al., 2003; Szameitat et al., 2002) and the maintenance of delayed intentions (Gilbert, 2011). Lateral PFC has also an important role in rule representation and the selection of correct rules (Brass & Cramon, 2002; Crone, Wendelken, Donohue, & Bunge, 2006), while the thalamus has also shown a role in cognitive flexibility during task switching (Rikhye, Gilra, & Halassa, 2018). Overall, this pattern reflects the cognitive demands of holding two tasks in mind, extending the results to the maintenance of social categorization sets.

In addition, we observed a set of task-negative regions such as mPFC, middle cingulate, OFC, IFG and temporal cortex which showed reduced signal when participants needed to hold an intended task set. Similarly, Landsiedel & Gilbert (2015) carried out an intention-offloading paradigm where participants had to remember a delayed intention, which they had to fulfil after a brief filled delay. During the maintenance of such intention they found a set of deactivated regions including mPFC, posterior cingulate cortex, infero-temporal cortex, and temporo-parietal cortex. Interestingly, these authors showed that when participants had the opportunity to offload intentions by setting an external reminder, this decrease of activation was ameliorated. This reduction suggests that these task-negative areas play a role in the representation of the delayed tasks. Pure blocks in our tasks are similar to the offloading condition in Landsiedel & Gilbert (2015), as participants needed to sustain only the initial task set to perform the correct social categorization. Therefore, taking these results into account, it seems unlikely that task-negative regions reflect simply a “default mode” (Fox et al., 2005), but rather that this deactivation is, to some extent, playing a functional role during task performance (Spreng, 2012).

Currently, multivariate pattern analyses are one of the most powerful approaches to study the information contained in different brain areas. In this work, we employed MVPA to study the regions representing initial and intended task sets. Importantly, a novelty of our approach was to do so by employing three different tasks instead of just two, unlike most previous studies (e.g. Haynes et al., 2007; Momennejad & Haynes, 2013). We were able to distinguish different regions representing initial and intended task sets. The currently-active initial task was decoded from left OFC, while information about the intended one was contained in the right IFG. Also, comparing between these two, a nearby OFC region together with the FG showed higher fidelity of the representation for the initial vs. intended task set. The left OFC region found for the initial task decoding has been previously associated with representations of facial information related to social categories (Freeman, Rule, Adams, & Ambady, 2010) and it also facilitates object recognition in lower level areas such as the FG (Bar et al., 2006). In addition, the FG showed greater fidelity of the representation of initial vs. intended task sets, which could indicate the allocation of attentional resources to process current relevant information in earlier perceptual regions. Besides having a prominent role in processing faces (Haxby et al., 2000), previous studies have been able to decode task-relevant information related to social categories (e.g. female vs. male, black vs. white) in perceptual regions such as the FG (Gilbert et al., 2012; Kaul et al., 2014, 2011; Stolier & Freeman, 2017). On top of this, previous work using facial stimuli in the field of social perception and prejudice has shown how the FG is influenced by stereotypical associations and evaluations of social categories associated with the OFC (Gilbert et al., 2012; Stolier & Freeman, 2016). Our data fit and extend this work, suggesting that both OFC and FG contain information about the appropriate task set to perform the initial classification, pointing out the role of both high and lower-level perceptual regions in the representation of ongoing task sets performed on facial stimuli.

Conversely, decoding of the intended task was possible from the IFG. Previous work has related lPFC with the representation of intentions (Haynes et al., 2007; Momennejad & Haynes, 2012; Momennejad & Haynes, 2013; Soon et al., 2008) and it has also been linked to task-set preparation (e.g. González-García, Arco, Palenciano, Ramírez, & Ruz, 2017; González-García, Mas, de Diego-Balaguer & Ruz, 2016; Sakai & Passingham, 2003). This result, however, contradicts the proposal of Gilbert (2011) in which lPFC would serve as a general store for delayed intentions without information about their content. Nonetheless, while Gilbert (2011) decoded simpler stimulus-response mappings, in our study we classified abstract task-set information, as previous studies that also decoded intention from lPFC did (e.g. Haynes et al., 2007; Momennejad & Haynes, 2013). Therefore, our work agrees with Momennejad & Haynes (2013), suggesting that lPFC may represent delayed intentions when their content is abstract enough.

Interestingly, we found a region in vmPFC/OFC that contained information about both initial and intended task sets. We examined if this vmPFC/OFC region contained overlapping representations of both initial and intended task sets by performing a cross-classification analysis, and observed a lack of generalization from the two sets of representations. Our cluster is close to the mPFC region found by previous intention studies (e.g. Gilbert, 2011), but located in a more ventral area. Notably, in our paradigm both the initial and intended tasks that participant performed were equally relevant and were based on the same set of stimuli. Altogether, this suggests that mPFC may be recruited when relevant task sets need to be held for a period of time, irrespective if this information is being using at the moment or later. This result also fits with a recent proposal (Schuck, Cai, Wilson, & Niv, 2016) characterising this region as a repositoiry for cognitive maps of task states. Specifically, the vmPFC/OFC would represent task-relevant states that are hard to discriminate based on sensory information alone (Schuck et al., 2016; Wilson, Takahashi, Schoenbaum, & Niv, 2014), similar to our study. Overall, this suggests a role for this region in the maintenance of task-relevant information, regardless of when it is needed. Given that cross-classification between initial and intended task sets was not possible in this region, this would be compatible with multiplexed representations of the two types of information (i.e. representations using orthogonal representational codes). It could also be compatible with nonoverlapping populations of cells representing initial and intended tasks within the voxels in this region. Further, it could be argued that the lack of cross-classification between initial and intended tasks could be explained by differential processes underlying the representation of these tasks. That is, the initial one could be represented simply by the activation of the S-R associations, while the intended one would rely on the maintenance of a general task set. Nonetheless, an explanation relying solely on this distinction seems unlikely, since we observed interference effects. Even though the interference effect for accuracy was smaller before the switch, for reaction times we observed interference only during this period. This indicates that the S-R association for both tasks is active prior the switch, although there could be some distinctions in the way these response representations are maintained. However, the lack of significant cross-classification could also reflect simply a lack of statistical power.

Despite our decoding results, we did not find the dissociation pattern that we predicted for initial and intended task sets. Although we could decode intended task sets from mPFC, consistent with previous studies (Gilbert, 2011; Haynes et al., 2007; Momennejad & Haynes, 2012; Momennejad & Haynes, 2013), we did not find a frontoparietal representation of initial task sets as previous studies have (Qiao et al., 2017; Waskom et al., 2014; Woolgar et al., 2011b). This discrepancy could be explained by differences in the stimuli and task employed. Previous studies have decoded task information in the form of classification between different stimulus-responses mappings or different types of stimuli. In contrast, in our task we employed the same stimuli for all tasks, and the differences between the information to decode relied in the representation of the social category needed for the specific part of the miniblock, rather than perceptual features (e.g. the colour or shape of target stimuli), which may be easier to decode (Bhandari, Gagne, & Badre, 2018).

To conclude, in the present work we examined the representation of A) currently-active initial task sets, and B) intended task sets during a social categorization dual-sequential task. Crucially, we directly examined the common and differential representation of initial and delayed task sets, extending previous work studying these mechanisms separately. Moreover, we employed faces as target stimuli to complement prior research. Apart from replicating previous findings in dual tasks with social stimuli, we show how task set information was contained in different regions, depending on when it was needed. Thus, currently-active initial tasks were represented in specialized regions related to face processing and social categorization such as the OFC and FG, while intended ones were represented in lPFC. On top of that, we showed a common brain region in vmPFC/OFC maintaining a general representation of task-relevant information, irrespective of when the task is performed, albeit it is not clear whether overlapping patterns of activation represent both types of information. Moreover, the results from the classification after the switch suggest that the representation of the two relevant tasks varies once the switch takes place and the initial task is no longer active. Future research should. further characterize the representational format of relevant task information depending on when it is needed and examine the structure of the task set representation within these regions, for instance using Representational Similarity Analysis (RSA, Kriegeskorte, Mur, & Bandettini, 2008). Also, studying the interaction of sustained task sets with specific conditions of each trial would extend our knowledge of the representational dynamics of current and intended task-relevant information. Last, due to the social significance of faces, one step forward would be to examine how their representation may vary in social scenarios, contexts in which faces and other social stimuli are particularly relevant to guide behaviour (Díaz-Gutiérrez, Alguacil, & Ruz, 2017).

## Funding

This work was supported by the Spanish Ministry of Science and Innovation (PSI2016-78236-P to M.R.) and the Spanish Ministry of Education, Culture and Sports (FPU2014/04272 to P.D.G.).

## Acknowledgments

This research is part of P.D.G’s activities for the Psychology Graduate Program of the University of Granada.

