## Supplementary Materials for "Neural representation of current and intended task sets during sequential judgements on human faces"

**fMRI non-parametric analyses**

Univariate

Permutations tests were carried out employing statistical non-parametric mapping (SnPM13, <http://warwick.ac.uk/snpm>) with 5000 permutations. We performed cluster-wise inference on the resulting voxels with a cluster-forming threshold at 0.001, which was later used to obtain significant clusters (FWE corrected at p<0.05).

Decoding

For the MVPA, we followed the method proposed by Stelzer, Chen, & Turner (2013), especially suitable for MVPA data. As in Arco, González-García, Díaz-Gutiérrez, Ramírez, & Ruz (2018), we permuted the labels and trained the classifier 100 times for each participant. After normalizing the resulting maps to an MNI space, for each participant, we randomly picked one of these maps, averaged them and obtained a map of group permuted accuracies. This method was repeated 50000 times to build an empirical chance distribution for each voxel. The 50^th^ greatest value corresponds to the threshold for statistical significance at p<0.05, FDR corrected.

**fMRI non-parametric results**

Univariate

*Pure vs. Mixed blocks*

Before the switch, the right middle frontal gyrus (k = 77; 42, 44, 23) showed higher activation when participants had to maintain two tasks versus one. Conversely, in this scenario, we observed decreased activation in a set of regions. Here, we observed cluster deactivation in bilateral middle cingulate cortex (k = 274; -3, -10, 33), bilateral medial prefrontal cortex (mPFC; k = 86; -6, 44, 41), left orbitofrontal cortex (OFC; k = 148; -33, 32, -16), right inferior frontal gyrus (IFG)/OFC (k = 193; 54, 32, -1), right parahippocampal/fusiform gyri and hippocampus (k = 37; 27, -19, -22), left parahippocampal/fusiform/inferior temporal gyri (k = 49; -21, -19, -28) and left lingual and parahippocampal gyri (k = 257; -9, 52, 5).

*Switch vs. non-switch trials*

Transient activity during switching tasks was observed in the bilateral brainstem and thalamus (k = 380; -3, -28, -22) as well as in a cluster including the left IFG/MFG and precentral gyrus (k = 210; -51, 11, 17). Further, these trials increased activation bilaterally in the anterior insula/IFG (k = 31; -24, 23, -7/k = 30; 24, 26, -4).

*Incompatibility effects*

Last, we assessed whether the compatibility effects observed on behaviour were matched at a neural level, by comparing incompatible vs. compatible trials before and after the switch. Data showed higher activation in the left IFG (k = 42; -48, 35, 20) for incompatible > compatible trials before the switch. The opposite comparison (compatible > incompatible before the switch) or incompatible vs. compatible contrasts after the switch did not yield any significant results.

Decoding

First, we averaged all individual classification maps to examine the regions sensitive to any kind of task (initial or intended) during the period prior to the switch. Here, we found that the rostromedial PFC/orbitofrontal cortex (OFC) presented significant accuracies above chance (k = 48; 15, 56, -10). Further, looking at the classification of the initial task only, we observed a cluster in the right OFC (k = 13, 15, 56, -13). However, in the classification for the intended task, we did not find any region showing significant accuracies above chance.

Last, when comparing decoding accuracies between the initial and intended tasks (subtracting initial – intended accuracy maps), we observed significantly higher accuracies for the representation of the initial (vs. intended) tasks in a cluster (k = 24; 39, -13, -25) covering the right FG and the hippocampus. Nonetheless, the opposite contrast (intended vs. initial) did not show any cluster with significant accuracies above chance.

**RT analyses between miniblock pairs and correlation with decoding accuracies**

In order to assess the influence of RT difference between tasks in decoding accuracies, we carried out a comparison between miniblock pairs. In line with the classification analyses, we performed paired t-tests between these pairs during the period before the switch: EG vs. RG/ER vs. GR/GE vs. RE for initial task differences and EG vs. ER/GE vs. GR/RG vs. RE for the intended task. Doing this, two contrasts of the initial tasks were significant: emotion vs. race (EG vs. RG, p=.003) and emotion vs. gender (ER vs. GR, p =.008), but no differences for gender vs. race (GE vs. RE, p>.1) or any of the comparisons for the intended tasks (all ps>.3).

Then, similarly to Crittenden et al. (2015) we obtained absolute RT differences for the initial task and intended task discriminations and conducted a Spearman's correlational analysis of RTs against decoding accuracies for each result ROI. As can be seen in Table 1, results show that decoding is not related to response times’ differences between the conditions that have been classified.

**Supplementary Table 1.** Spearman’s correlational analysis of RTs against decoding accuracies.

|  | Spearman r | p-value |
| --- | --- | --- |
| **Relationship between the general decoding ROI and response times.** |  | |
| general RT differences and general decoding | -.181 | .134 |
| initial RT differences and general decoding | -.201 | .322 |
| intended RT differences and general decoding | -.141 | .442 |
| initial RT differences and general decoding, initial | -.196 | .442 |
| intended RT differences and general decoding, intended | -.252 | .164 |
| **Relationship between the initial decoding ROI and**  **response times** |  | |
| initial RT differences and OFC decoding | .008 | .965 |
| **Relationship between the initial > intended decoding ROIs and response times** |  | |
| initial RT differences and OFC decoding | .087 | .636 |
| initial RT differences and FG decoding | .025 | .892 |
| **Relationship between the intended decoding ROI and response times.** |  | |
| intended RT differences and IFG decoding | .152 | .407 |
